# How well do RNA-Seq differential gene expression tools perform in a eukaryote with a complex transcriptome?

**DOI:** 10.1101/090753

**Authors:** Kimon Froussios, Nick J. Schurch, Katarzyna Mackinnon, Marek Gierliński, Céline Duc, Gordon G. Simpson, Geoffrey J. Barton

## Abstract

RNA-seq experiments are usually carried out in three or fewer replicates. In order to work well with so few samples, Differential Gene Expression (DGE) tools typically assume the form of the underlying distribution of gene expression. A recent highly replicated study revealed that RNA-seq gene expression measurements in yeast are best represented as being drawn from an underlying negative binomial distribution. In this paper, the statistical properties of gene expression in the higher eukaryote *Arabidopsis thaliana* are shown to be essentially identical to those from yeast despite the large increase in the size and complexity of the transcriptome: Gene expression measurements from this model plant species are consistent with being drawn from an underlying negative binomial or log-normal distribution and the false positive rate performance of nine widely used DGE tools is not strongly affected by the additional size and complexity of the *A. thaliana* transcriptome. For RNA-seq data, we therefore recommend the use of DGE tools that are based on the negative binomial distribution.

## Introduction

Short read RNA sequencing (RNA-seq) has become the method of choice for transcriptome-wide quantification of gene expression and the analysis of differential gene expression between experimental conditions (Mortazavi et al. 2008;Nagalakshmi et al. 2010). RNA-seq data analysis typically involves aligning short sequence fragments (reads) to a reference genome or transcriptome, counting the resulting alignments that fall within an annotated feature region, then identifying any significant differences between two conditions. More than a dozen computational tools have been developed to identify Differential Expression (DE) from RNA-seq data and each makes assumptions about the variability of the RNA-seq expression measurements across replicates (Anders and Huber 2010;Hardcastle and Kelly 2010;Robinson et al. 2010;Wang et al. 2010;Tarazona et al. 2011;Li et al. 2012;Lund et al. 2012;Trapnell et al. 2012;Li and Tibshirani 2013;Leng et al. 2013;Frazee et al. 2014;Law et al. 2014;Love et al. 2014;Moulos and Hatzis 2014;Ritchie et al. 2015). Based on these assumptions, the tools calculate the probability that two sets of measurements come from the same statistical distribution, thus determining whether a genuine shift in expression is a more likely explanation for the observed values than random chance. Assuming an incorrect distribution can lead to poor False Discovery Rate (FDR) control and inaccurate True Positive (TP) identification in the DGE calls. Such errors will propagate downstream into the biological interpretation of the DE results. Most commonly, these tools are used to identify DE for genes (Differential Gene Expression - DGE), however they are increasingly being used to identify DE of other genic regions (i.e. exons, spliced transcripts, etc.) (Wood et al. 2013;Frazee et al. 2014;Gaidatzis et al. 2015).

Several studies have assessed the performance of DGE tools (Busby et al. 2013;Soneson 2014;Bullard et al. 2010;Rapaport et al. 2013;Seyednasrollah et al. 2015;Li et al. 2012;Lund et al. 2012;Trapnell et al. 2012;Li and Tibshirani 2013;Leng et al. 2013;Frazee et al. 2014;Law et al. 2014;Love et al. 2014;Moulos and Hatzis 2014;Ritchie et al. 2015). However, these studies were carried out using either simulated data or biological data that was originally designed for a different purpose. Although a few of these studies have explored high biological replication by leveraging publicly available datasets on individuals within a species (Seyednasrollah et al. 2015;Guo et al. 2013;Soneson and Delorenzi 2013;Burden et al. 2014;Bottomly et al. 2011), most have a limited level of replication. Recently, a study was performed in yeast specifically to test both the underlying statistical properties of RNA-seq data across biological and technical replicates and the influence of replication on DGE results (Gierlinski et al. 2015;Schurch et al. 2016). With 48 biological replicates per condition, the distribution of read counts per gene was found to be most consistent with a negative binomial (NB) distribution, and the FDR of many DGE tools was demonstrated to be low at all replication levels. However, the number of replicates was shown to affect the tools’ sensitivity for identifying differential expression, particularly at low fold-changes. The authors recommended that future RNA-seq experiments that focus on identifying DE have at least six replicates per condition in order to reliably detect genes with fold-changes greater than a factor of 2, rising to at least twelve replicates to reliably detect the majority of differentially expressed genes regardless of fold-change.

In this paper, RNA-seq data from 17 wild-type biological replicates of *Arabidopsis thaliana* were used to explore read count measurements across replicates and the FDR of DGE tools. Although *A. thaliana* has a relatively small genome, its transcriptome is similar in scale and complexity to that of model mammal species (Arabidopsis Genome Initiative 2000;Carvalho et al. 2013;Krishnakumar et al. 2014) and its genome is extensively annotated. Accordingly, conclusions from the high replicate RNA-seq study presented here should provide useful guidance for work in other complex eukaryotes as well.

## Results

### Consistency among replicates

Our dataset consisted of 100-base reads from 17 wild type *Arabidopsis thaliana* samples with sequencing throughput of at least 77 x 10^6^ read pairs per sample. The samples were collated from three separate experiments (see Materials and Methods for details). The global gene expression measurements from 16 of the 17 wild-type biological replicates are well correlated, irrespective of the different experiments (R>0.99, Figure 1). Replicate 11 correlates less well with all the other replicates (0.83 *≤ R ≤* 0.87, Figure 1) due to less efficient removal of ribosomal RNA during sample preparation, as evidenced by higher read counts across ribosomal genes by at least an order of magnitude compared to the other replicates (Supplementary Table 1) and high level of multi-mapping reads (Supplementary Table 2). Consequently, replicate 11 was excluded from the subsequent analyses. Additionally, a low but uniform level of read coverage across the genome was observed in the replicates from ExpB (replicates 8-14), explaining the marginally lower correlation between the replicates of this experiment and the other replicates (Figure 1, right panel).

**Figure 1.**
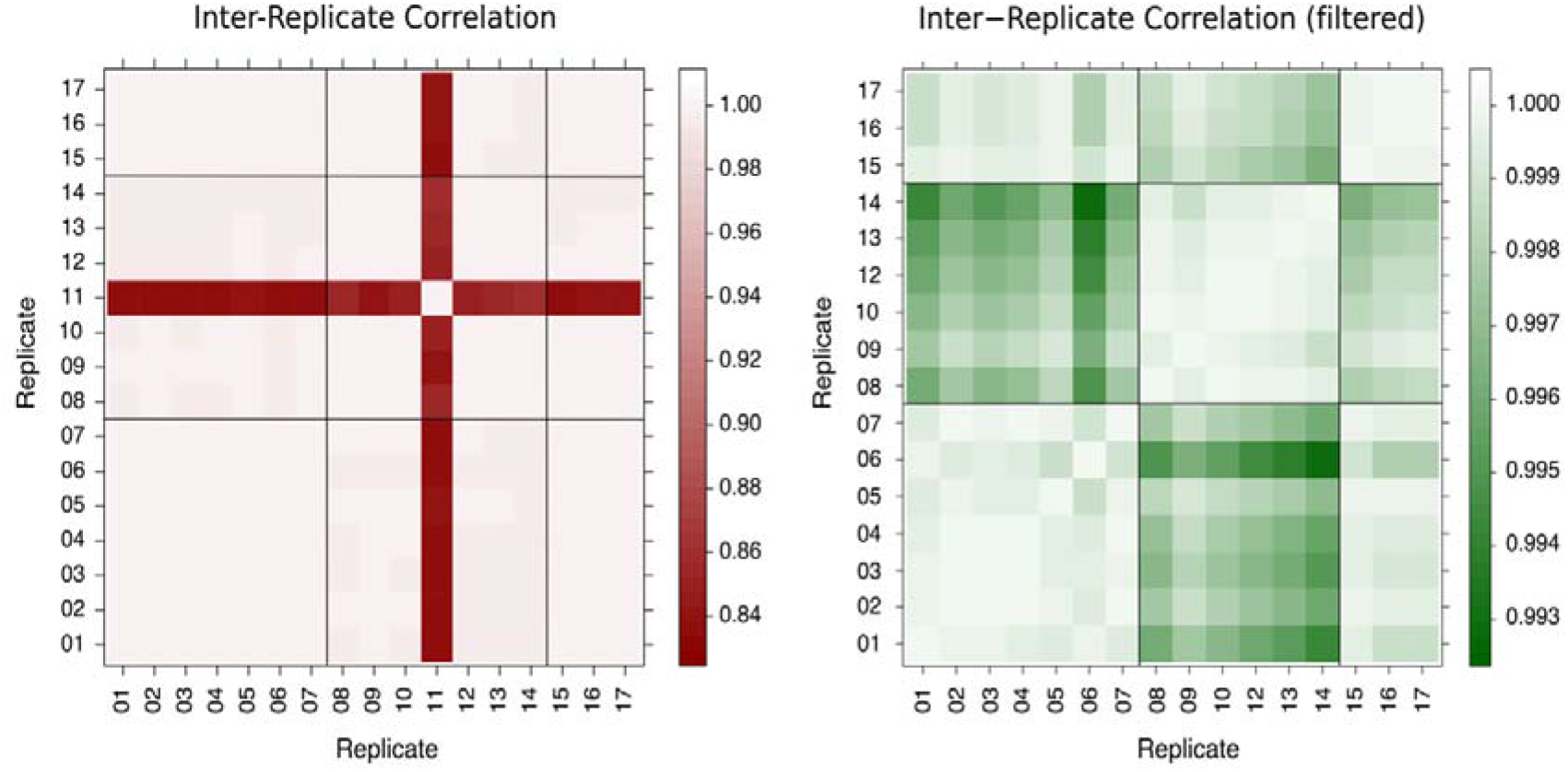
[Left]: Pairwise Pearson correlation of gene expression for all 17 replicates. Apart from replicate 11, all replicates correlate very well. [Right]: Same as left, but with replicate 11 filtered out, allowing the patterns of correlation among the remaining 16 replicates to be better seen.

### Distribution of gene read counts across replicates

Figure 2 shows that the distribution of expression measurements across replicates is consistent with being drawn from a negative binomial distribution for 100% of the genes, and is consistent with a log-normal distribution for 98% of genes. In contrast, 23% of genes have expression distributions that are not consistent with a normal distribution and more than 70% are inconsistent with a Poisson distribution.

**Figure 2.**
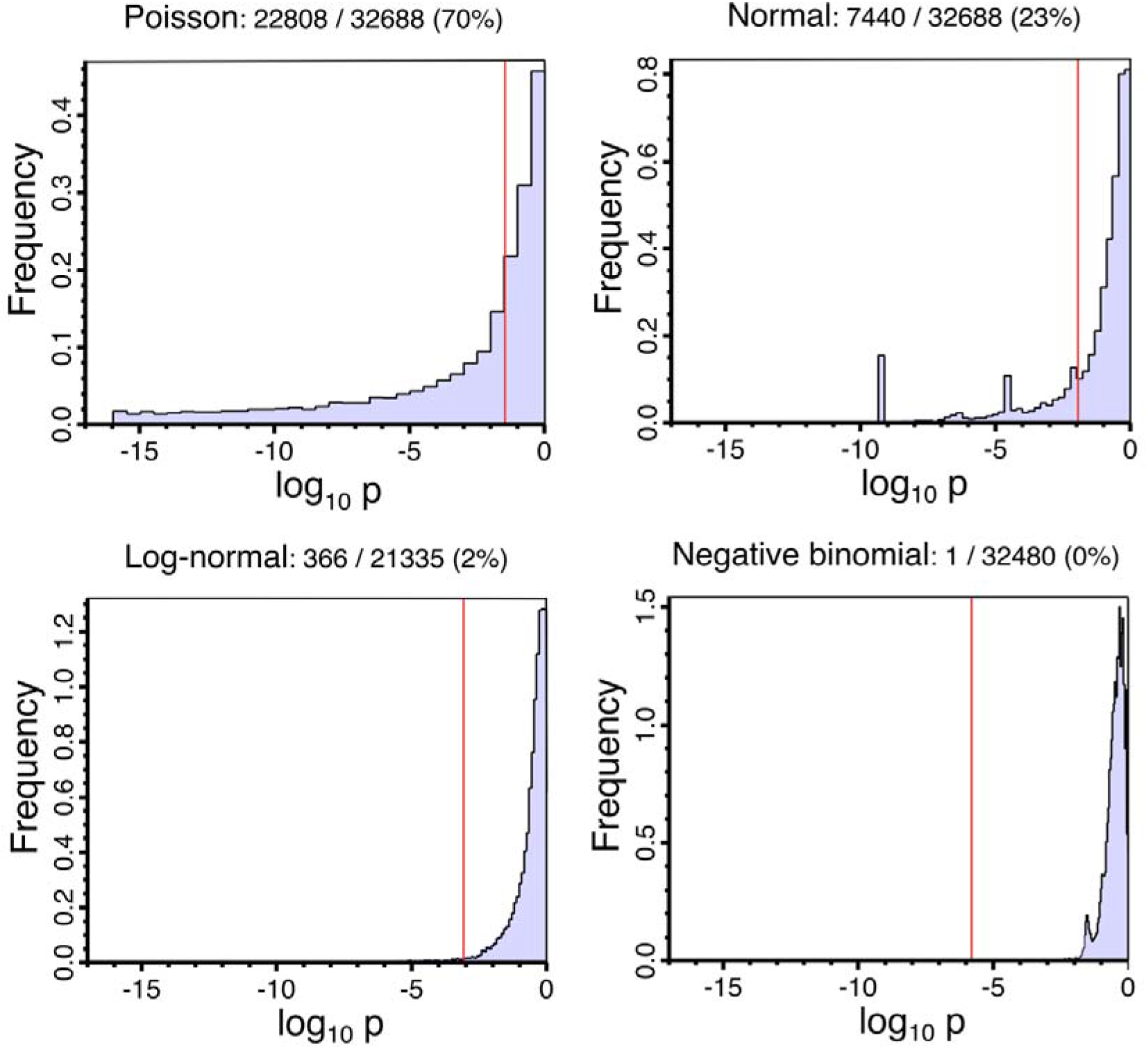
Histograms of the probability that the genes’ fragment counts across replicates are compatible with each of the four specified distributions. The fraction of genes rejecting the distribution model is given above each plot.

As mentioned previously, replicates 8-14 presented a low level of uniform read coverage along the genome. We believe this to be noise consistent with a small amount of DNA contamination in the affected samples. Such reads have the potential to interfere with the fitting of statistical distributions to the data, as they can make silent genes artificially appear as expressed. Indeed, Figure 3 shows that approximately 6,000 of the genes annotated in Araport11 appear to be lowly expressed in the affected replicates, but are not detected in the ten replicates from the other two experiments (replicates 1-7 and 15-17). The potential for this noise to impact on the distribution measurements was assessed by comparing the fit of the distributions to the full dataset of 16 replicates against the the fit without the noisy replicates of ExpB. As a control, the distribution fitting was also performed with the exclusion of replicates 1-7 in place of the noisy ones. In both cases, the fit of the distributions to the reduced dataset was very similar to that of the full dataset, with the exception of the normal distribution where, in both the noise exclusion and the control, the fraction of the gene data that is inconsistent with the distribution was reduced from 23% to ~10%. The lack of difference between excluding the noisy replicates and excluding the control replicates demonstrates that the apparent improvement of model fit is due to the reduction of statistical power caused by the smaller number of replicates, rather than to a cleaner signal.

**Figure 3.**
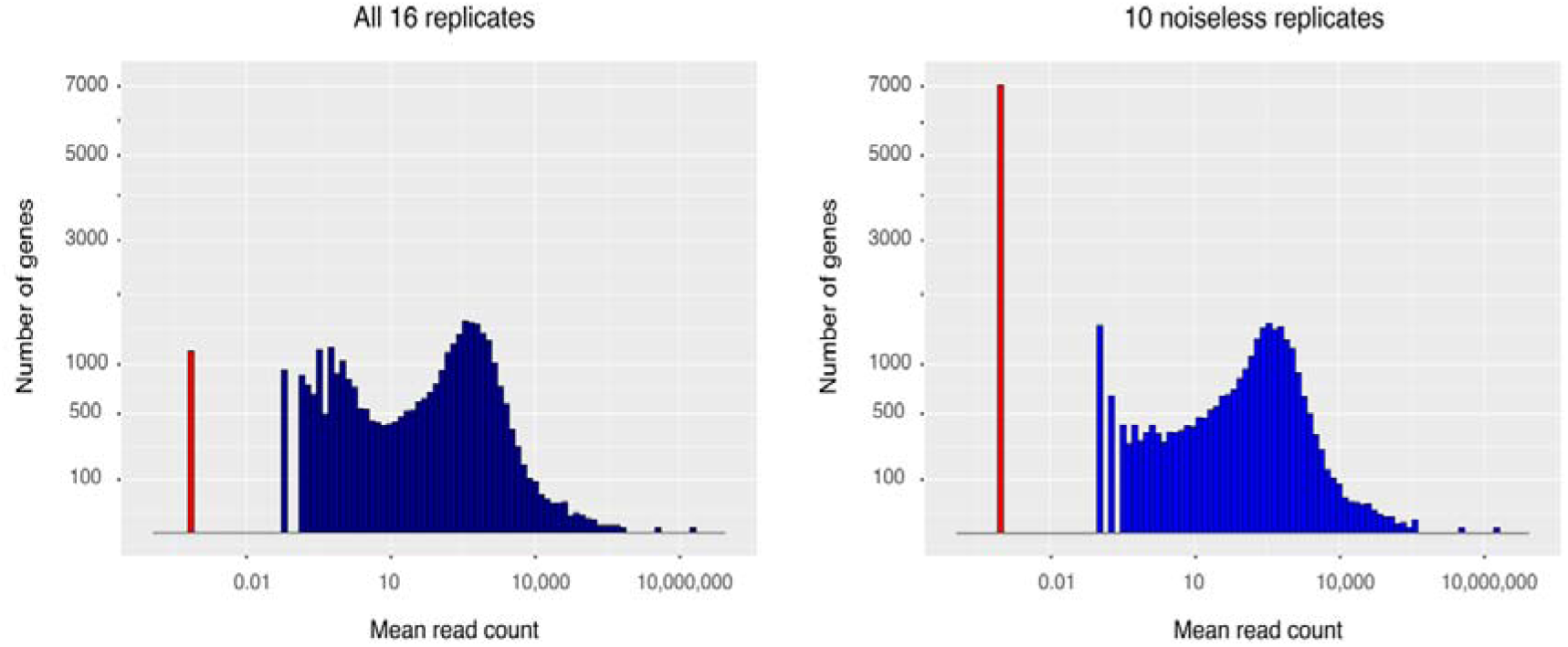
Distribution of gene expression. Each gene is represented by the mean of its read count estimates across replicates. The various levels of non-zero expression are shown in blue. Because the x-axis is in logarithmic scale, the number of genes with zero expression was added manually at an arbitrary but distinct location on the axis and is shown in red.

### False positive behavior of differential gene expression tools

The distribution of the false positive fraction as a function of the number of replicates, for each differential expression tool, is shown in Figure 4. Most tools consistently control their FP fraction well at all numbers of replicates despite the presence of a small number of outlier results. *DEGseq* fails to control its FP fraction adequately, likely due to over-estimation of the performance is worse than the other tools at all the tested numbers of replicates, suggesting that it is a poor choice for calling SDE.

**Figure 4.**
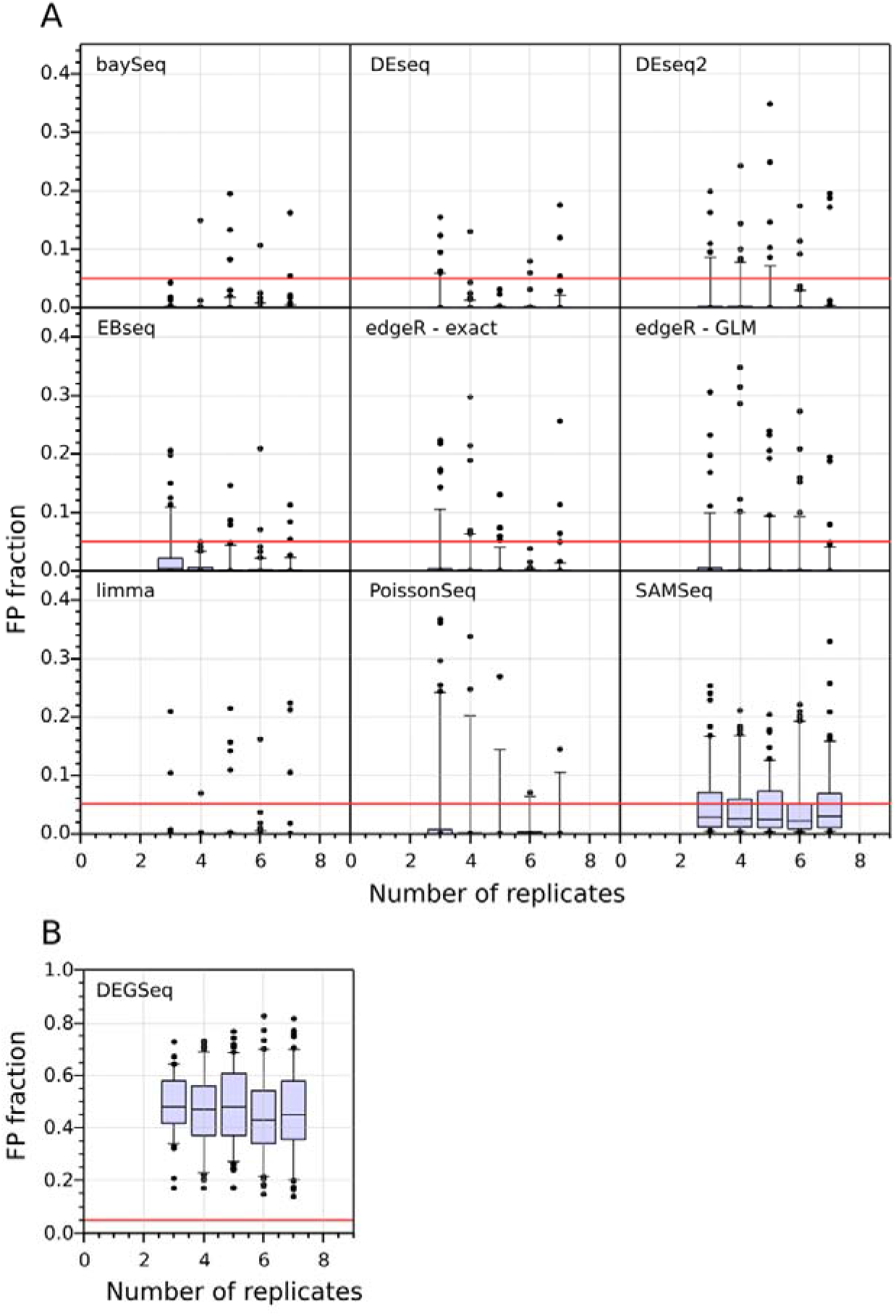
False Positive Rates in WT vs. WT comparisons of Differential Gene Expression, measured across 100 bootstrap iterations and across a range of sample sizes from 2 to 7 replicates per condition. Panel [A] and [B] differ on the range of the Y-axis. DEGSeq displays poor FPR performance (nearly 50% of its positives are false). The median FPR for all the other tools is below 5%. The performance of the tools is connected to their choice of statistical model (Table 1), with the lowest FPR tools using the negative binomial or lognormal distributions.

**Table 1.** RNA-seq differential gene expression tools used in this study.

| Name | Assumed Distribution | Normalization | Description | Version | Citations<sup>1</sup> |
| --- | --- | --- | --- | --- | --- |
| *baySeq* (Hardcastle and Kelly 2010) | negative binomial | Internal | Empirical Bayesian estimate of posterior likelihood | 2.4 | 186 |
| *DEGseq* (Wang et al. 2010) | binomial | None | random sampling model using Fisher's exact test and the likelihood ratio test | 1.24.0 | 411 |
| *DESeq* (Anders and Huber 2010) | negative binomial | DEseq | Shrinkage variance | 1.22.0 | 2384 |
| *DESeq2* (Love et al. 2014) | negative binomial | DEseq | Shrinkage variance | 1.10.0 | 590 |
| *EBSeq* (Leng et al. 2013) | negative binomial | DEseq (median) | Empirical Bayesian estimate of posterior likelihood | 1.10.0 | 120 |

<sup>1</sup> as reported by PubMed Central articles that reference the listed source on the 23<sup>rd</sup> Aug 2016.

|  |  |  |  |  |  |
| --- | --- | --- | --- | --- | --- |
| *edgeR*<br>(Robinson et al. 2010) | negative<br>binomial | TMM<sup>2</sup> | Empirical Bayes estimation &<br>either an exact test analogous to<br>Fisher's exact test but adapted<br>to over-dispersed data or a<br>generalized linear model | 3.12 | 2068 |
| *Limma*<br>(Ritchie et al. 2015) | Log-normal | TMM<sup>2</sup> | Generalised linear model | 3.26.2 | 266 |
| *PoissonSeq*<br>(Li et al. 2012) | Poisson log-<br>linear model | Internal | Score statistic | 1.1.2 | 48 |
| *SAMSeq*<br>(Li and Tibshirani 2013) | None | Internal | Mann-Whitney test with<br>Poisson resampling | 2.0 | 73 |

## Discussion

In this study the statistical assumptions made by tools that identify differential gene expression from RNA-seq read count data were validated in a high-replicate experiment. The findings show that the negative binomial (NB) and log-normal distributions are both good choices as models for the cross-replicate variability of RNA-seq read counts. The study demonstrates that 6 out of the 9 DGE tools examined here control their identification of false positives well even with only 3 replicates. In contrast, tools that assume distributions other than negative binomial or log-normal showed higher FPR, at all replication levels. These results reinforce similar conclusions previously reached by studies of the yeast transcriptome (Gierliński et al. 2015;Schurch et al. 2016).

The transcriptome of *A. thaliana* is considerably more complex than *S. cerevisiae,* with almost four times the number of protein-coding genes (27,667 in *A. thaliana,* 7,126 in *S. cerevisiae)* and widespread alternative splicing and alternative polyadenylation. The similarity of the results from these two diverse organisms suggests that the conclusions of both studies are applicable to a wide range of eukaryotes.

However, the concept of gene expression in complex transcriptomes is confounded by the presence of alternative transcript isoforms, which give the organism additional means to regulate a gene’s expression. This type of regulation is not necessarily reflected in changes to the total transcriptional output by a gene. Ideally, expression studies should aim to quantify the abundance of alternative isoforms individually and independently. Interestingly, the sum of independent random variables with a NB distribution itself has a NB distribution. Thus, the conclusion that a NB is a suitable model for gene expression variability across replicates is consistent with the hypothesis that the underlying variability of expression of the individual isoforms also follows the NB distribution. This suggests that tools originally intended for the study of differential gene expression may also be appropriate for studying differential transcript expression (DTE).

In summary, our analyses show that the statistical properties of gene expression are similar between simple and complex eukaryotes, and validate the model assumptions of the best-performing DGE tools, thus justifying their use in a broad range of organisms.

## Materials and Methods

### Sample Preparation & Sequencing

The RNA-seq data for this study are wild-type (WT) *Arabidopsis thaliana* Colombia-0 (Col-0) biological replicates from three separate experiments (hereafter ExpA, ExpB & ExpC). Briefly, for all three experiments WT *A. thaliana* Col-0 seeds were sown aseptically on MS10 plates. The seeds were stratified for 2 days at 4^o^C and then grown at a constant 21^o^C under a 16-h light/8-h dark cycle for a further 14 days, at the end of which the seedlings were harvested. Total RNA was isolated from the seedlings with the RNeasy Plant Mini Kit (Qiagen) and treated with TURBO™ DNase (Ambion). 4 μl of ERCC spike-ins (External RNA Controls Consortium 2005) at a 1:100 dilution were added to 1 μg/6 μl of total RNA. Libraries were prepared using the Illumina TruSeq Stranded Total RNA with Ribo-Zero Plant kit. The libraries were sequenced on a HiSeq2000 at the Genomic Sequencing Unit of the University of Dundee. Two of the experiments, ExpA & ExpB, have 7 biological WT replicates (replicates 1-7 and 8-14 respectively) while ExpC has 3 (replicates 15-17), for a total of 17 biological WT replicates and ~1.7 × 10^9^ 100-bp paired-end reads across the three experiments. The plants were sown, grown, harvested and the libraries were prepared by the same lab, and the sequencing was performed on the same machine by the same people at the same sequencing facility and all the samples include the ERCC spike-ins which can verify the WT samples are consistent and comparable across experiments.

### Quality Control, Alignment & Quantification

The quality of the data was quantified using *FastQC* v0.11.2 (http://www.bioinformatics.babraham.ac.uk/projects/) with all the replicates performing as expected for high quality RNA-seq data with excellent median per-base quality (≥38) across >90% of the read length. The read data for each sample were aligned to the TAIR10 *A. thaliana* genome using the splice-aware aligner *STAR* v2.5.0a (Dobin et al. 2012). The index was built with `“--sjdbOverhang 99”` and the alignment was run with parameters: `“--outSAMstrandField intronMotif -- outSJfilterIntronMaxVsReadN 5000 10000 15000 20000 -- outFilterType BySJout -- outFilterMultimapNmax 2 -- outFilterMismatchNmax 5”`.

Read counts per gene were then quantified from these alignments with *featureCounts* v1.4.6-p4 (Liao et al. 2014) using the publically available Araport11 annotation (pre-release 3, 12/2015, comprising 33,851 genes) (Krishnakumar et al. 2014) with the parameters: `“-texon-ggene_id-s2-p-P”.`

These read counts were used without further processing to examine the FPR performance of nine DGE tools, allowing each tool to carry out its default normalization. The tools were used in the R v3.2.2 environment and installed through Bioconductor v3.2.

For the purposes of comparing the expression distribution models, consistently normalized data was required. As some of the distributions in question are discrete, normalized integer read counts were used for this purpose, which were calculated by randomly down-sampling read-pairs from each replicate to the level of the replicate with the lowest read depth. In the present study, the focus is on the collective behavior of gene expression, rather than the biological interpretation of the expression of any specific gene, so this type of normalization is appropriate here. However, it is not recommended for typical gene expression analysis studies, as some low expression signals can randomly be lost during resampling. After the normalization, each replicate consisted of ~77 x 10^6^ read pairs, which were then aligned to the genome and quantified as described above.

### Performing the tests

The read counts of each gene were tested against four theoretical distributions across replicates: normal, log-normal, Poisson and negative binomial (NB). For the normal and lognormal distributions the goodness of fit was determined using the test for normality from D’Agostino *et. al.* (1990). For the NB distribution, the method described by Meintanis (2005)was employed and for the Poisson distribution a *χ^2^* test (Fisher 1950). In each case, rejection of the null hypothesis was based on a Benjamini-Hochberg corrected critical p-value of 0.05 (Benjamini and Hochberg 1995).

In order to test the false positive rate (FPR) of the DGE tools, two sets of *n_r_* replicates were randomly selected without replacement from the pool of 16 WT replicates. Differential gene expression was then called on each of the set pairs with each of nine DGE tools (Table 1), allowing each tool to apply its default normalization. Since all sets are drawn from the same WT pool, every gene identified as significantly differentially expressed (SDE) is, by definition, a false positive. This process was repeated 100 times for each sample size in the range 2 ≤ *n_r_* ≤ 7 for each tool.

## Availability

The raw data for the 17 WT Arabidopsis thaliana datasets is available from the European Nucleotide Archive ([database accession to be inserted here]).

## Acknowledgements

The authors wish to thank Dr. Christian Cole and Dr. Pieta Schofield for useful discussions. The authors also thank the BBSRC for funding this research (BB/H002286/1; BB/J00247X/1; BB/M010066/1; BB/M004155/1).

